# Limitations and Optimizations of Cellular Lineages Tracking

**DOI:** 10.1101/2023.03.15.532767

**Authors:** N. Leibovich, S. Goyal

## Abstract

Tracking cellular lineages using barcodes provides insights across biology and has become an important tool. However, barcoding strategies remain ad-hoc. We show that elevating barcode insertion probability, and thus increasing the average number of barcodes within the cells, adds to the number of traceable lineages but decreases the accuracy of lineages’ inference due to reading errors. We discuss how this tradeoff informs optimal experimental design under different constraints and limitations. In particular, we explore the trade-off between accuracy and the number of traceable lineages, concerning limited resources, the cells and barcode pool features, and the dropout probability.

## I. INTRODUCTION

Cellular barcoding is a technique in which individual cells of interest are tagged with heritable identifiers called barcodes. Some barcoding techniques are based on unique insertion sites, yet commonly nowadays cells are labeled with unique nucleic acid sequences that can be tracked through space and time, and thus gain insights into cellular behavior [1, 2]. For example, it has been used to study lineages of T-cells [3], hematopoietic stem cells [4, 5], clonal dynamics of cancer cells [6–8], and mapping axonal projections [9] to name only a few. Tracking of cell lineages over time requires the insertion of a unique set of barcodes into each cell, propagating the cell population over time, and accurate reading of all cells’ barcode sets. Ideally, it allows us to detect clones throughout the observation times and associate them with their lineages.

The insertion of barcodes into cells may be carried out by various procedures [10–13]. For example, by manually assigning individual barcodes to cells one-by-one and thus guaranteeing unique barcodes to each cell [10, 11]. However, this method is limited to a small population of cells and barcodes. Currently, the most common, robust, and efficient method to barcode individual cells relies on the production of a large pool of barcoded vectors (e.g., viruses) which deliver the barcodes into the cells [12]. Importantly, by infecting the ensemble of cells with a pool of barcoded viruses, the actual number of viruses or bacteria that will enter any given cell is a stochastic process: some cells may absorb more than one infectious agent while others may not absorb any [4, 5, 14]. Thus, this method, inherently, produces non-injective barcode-cell matching, meaning a given barcode can be inserted into more than one cell, and some cells may have more than one barcode. Yet, for a sufficient complexity of the barcode library, each cell’s set of labeling barcodes is presumably unique [15].

The barcoding procedure allows for the identification of many cells’ lineages via reading their barcodes using bulk sequencing approaches or the single-cell sequencing method which examines the nucleotide sequence information from individual cells [2, 16]. Each method of sequencing; either the genome or RNA sequencing, presents some advantages alongside some challenges concerning their preparation, amplification, extraction, reading, and data analyzing techniques [17–19]. For example, singlecell RNA sequencing provides the expression profiles of individual cells. This presents an issue because some barcodes may be expressed at a low level, leading to ‘dropouts’ where the barcode is not read for the cell [20–23]. Additionally, failure to detect barcode sequences when using transcriptomic data may be due to epigenetic silencing over time, especially with cell fate conversions [24].

As mentioned, the randomness in the barcoding procedure may induce overlap between barcodes’ set within cells. Hence, the dropouts together with the barcodes’ set overlap may lead to errors, and thus to inaccurate identification of lineages. Note, that misidentifying clones due to dropouts may emerge even when every clone is infected by its unique set of barcodes. For example, in Fig. 1 we present some clones that were infected with unique barcode sets at some initial time. Then, after some propagation time when clones are re-observed, one aims to associate later measured clones with their ancestors and to gain insight into the clonal dynamics in some space, which can be either real-physical space such as neuronal, or gene-expression space. Note that some clones may have more than one barcode, and some clones might not have been infected at all. Consequently, some clonal detection errors and thus wrong lineage identification might arise. For example, some observed clones in a later time, have not been measured in earlier times, and some cells might be wrongly associated with other lineages, see Fig. 1.

**FIG. 1.**
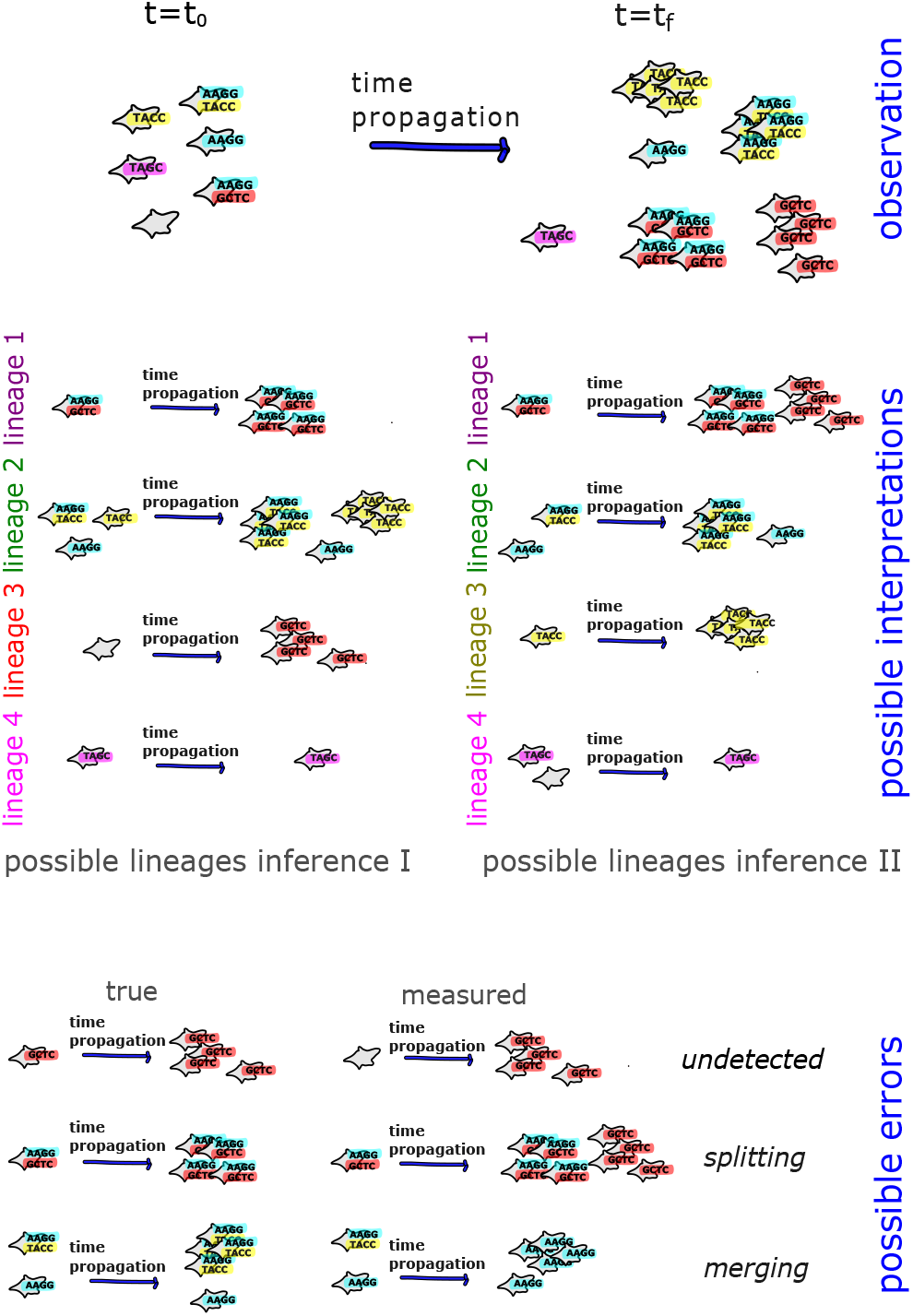
Single-cell barcoding allows tracking cell lineages with space and time. However, dropouts of barcodes throughout the observation complicate this task, even when the seeded barcodes’ sets are unique (upper panel). The presence of dropouts gives rise to several lineage-structure interpretations that can be inferred from the measured barcoded cells. In the above illustration, we present only two possibilities of lineages inference, although other lineages inferences are possible as well (middle panel). Wrongly or unidentified lineages may be occurred due to unmeasured barcodes in one (or more) snapshots, associating two lineages as a single lineage, or identifying a single lineage as two separate ones (lower panel).

In the random insertion method, namely the pool-of-barcodes procedure, one of the tunable parameters in the insertion setup is the multiplicity of infection (MOI), which is assumed to be the ratio between the number of viruses particles to the number of target cells present [25, 26]. Under the assumption that all virions are similarly infectious and all cells are similarly susceptible, the number of barcodes inserted into each cell presents Poisson distribution, and the average size of the barcodes’ set in each cell is exactly the MOI itself. We note that deviation from that assumption is empirically found [9, 27, 28], and we further discuss it below. Regardless of the distribution of barcodes’ sets size, we still consider cases of random insertion of barcodes, where barcodes may be inserted into more than one cell.

To overcome the lineages identification problem that emerges from barcodes dropouts, the experiment is usually designed such that the MOI is sufficiently low, and the barcodes library is highly diverse [14], see also Table I. Low MOI aims to reduce the number of clones with more than one barcode. A highly complex barcode library decreases the probability of any overlap between sets of barcodes. However, using too low MOI or a too diverse barcode pool may be excessive or unnecessary considering the desired accuracy of the empirical data, and the overall goal of the experiment [18].

**TABLE I.** Cellular barcoding parameters in a variety of systems.

| system [Ref.] | #cells | #barcodes | MOI | % labeled cells |
| --- | --- | --- | --- | --- |
| embryonic data [14] | $7.4 \cdot 10^4$ | $\sim 10^6$ | 0.2 (used)<br>0.15 (measured) | $\sim 20\%$ |
| hematopoietic stem cells [5] | | | 1 | $\sim 85\%$ |
| clonal dynamics [27] | $2 \cdot 10^6$ | 20,000 | 0.05 – 0.1 | 5% – 10% |
| neuronal structure [9] | $\sim 10^8$ (theory)<br>$\sim 10^2$ (exper.) | $\sim 10^{18}$ (theory)<br>$\sim 10^6$ (exper.) | 0.43 <sup>a</sup> | $\sim 80\%$ |
| induced pluripotent stem cells [30] | 170,000 –<br>230,000 | 50K –<br>16,000K | 0.35 –<br>0.89 | 29.1% –<br>59.1% |
| patient-derived xenografts [31]<br>of breast cancer | | | 0.07 <sup>b</sup><br>0.13 <sup>b</sup><br>0.23 <sup>b</sup> | $7 \pm 4.99\%$ <sup>c</sup><br>$12.18 \pm 8.13\%$ <sup>c</sup><br>$20.35 \pm 7.71\%$ <sup>c</sup> |
| clonal dynamics in reprogramming [28] |  | 36,000 | 0.05 – 0.1 | 5% – 10% |
| cells fate [32] | 5,000 – 10,000 | $0.5 \cdot 10^6$ | | |
| synaptic networks [33] | | $1.29 \cdot 10^6$ | $\sim 0.15$ <sup>d</sup><br>$\sim 1.5$ <sup>d</sup><br>$\sim 15$ <sup>d</sup> | $\sim 8.57\%$ <sup>e</sup><br>$\sim 32.81\%$ <sup>e</sup><br>$\sim 44.44\%$ <sup>e</sup> |

<sup>a</sup> infected neurons contained $1.2 \pm 0.1$ barcodes each [9]
<sup>b</sup> we estimated it using the assumption in [31] that it follows Poisson statistics
<sup>c</sup> mean ± STD, see [31]
<sup>d</sup> unique barcodes mean ± STD for neurons/astrocytes: MOI $\sim 0.15 : 1.07 \pm 0.009 / 1.03 \pm 0.01$ . MOI $\sim 1.5 : 1.42 \pm 0.01 / 1.14 \pm 0.01$ . MOI $\sim 15 : 1.6 \pm 0.01 / 1.2 \pm 0.02$ , see [33]
<sup>e</sup> we calculated it using the absolute number of infected and uninfected cells given in [33]

In the following, we examine the lineages’ identification quality, with respect to the MOI and the barcode pool complexity, subject to the bounded resources inherently emerging in any experimental setup. Excessive inflation of the barcodes’ library or the number of cells may be found redundant concerning the desired goal. Here we show that a certain barcode library complexity, together with some specific range of MOI, might be sufficient for complying with the observation purposes. To do so, we use synthetic data, simulated from a stochastic model. We emphasize though, that we consider and model only the dropout effect; which means where a barcode may be either measured or dropped. Other empirical information or measurement errors, such as barcode swapping [29], are left for further research.

## II. MODEL

We consider an ensemble of cells with a total of *S* cells at initial time *t* = 0. Then, we randomize an ‘insertion’ of barcodes from a pre-prepared pool with *B* different barcodes into the ensemble of cells. We note that *B* refers to the diversity or complexity of the barcodes’ pool, which means the number of *unique* barcodes in the pool. Note that *B* is different from the total number of barcode units.

The insertion procedure may be considered binomial or skewed, corresponding to whether the barcodes’ frequency within the pool is uniform or not, as well as to the susceptibility of the cells. After the initialization step, each cell may be associated with no barcodes (and thus is not observed), or with a combination of several barcodes. Note that, as mentioned, due to the randomness in the insertion process, each barcode may be inserted into more than one cell, and the set of integrated barcodes for each cell is not guaranteed to be unique. In the Supplementary Information (SI) App. A we show simulation results in agreement with the analytical prediction; uniform barcodes’ frequencies and cells’ susceptibility yields binomial distributions for both the number of cells with a given barcode as well as the number of barcodes integrated into a cell.

To pose the problem of lineages tracking, we propagate in time the population of the barcoded cells. In each generation, the entire cell population is multiplied and sampled such that the population size remains constant after each time step [34, 35]. As a result, the number of clones in each passaging decreases, while the mean clone size increases. In the simulation presented in this document we propagate the system for a total of 15 generations, but keep for analysis only the clone populations in the seeding stage, and of generations 5, 10, and 15; namely, we record and analyze only these four times frames. We note that most lineages are lost through the passaging events even when no dropouts occur, see additional simulation results in SI App. B

Note that here we have considered neutral dynamics across all cells, which means all cells proliferate at the same rate such that no clone possesses an advantageous growth rate over the others. However, that assumption might be different depending on the context, for example, it has been shown in reprogramming populations that some cells reprogram much faster thus the propagation would be biased in such a scenario [28].

We also drop some pre-inserted barcodes from the tracked cells to include the dropout effect. The dropout probability is assumed to be uniform resulting in a binomial distribution. Hence, for all propagated cells with their corresponding labeling barcode sets, there is a probability to drop some barcodes, which means not reading them. This probability is considered constant and thus independent of the integrated barcodes’ set size, the barcode, or the cells’ identity. Further details about the model, including the insertion, dropouts, and propagation, are given in SI Apps. A and B.

## III. MAIN RESULTS

For given measurement limitations, which correspond to dropout probability, the main strategies used for improving the precision of the lineages inference in a given experimental setup, are to combine both increasing the barcodes’ library size and decreasing the MOI. Indeed, increasing the barcodes’ pool complexity reduces the probability that any specific barcode is integrated into two different cells, thus the overlap between barcodes’ sets is very low. However, beyond some barcode pool complexity, the barcodes’ sets’ overlapping probability is nearly zero thus increasing the complexity even further is not beneficial. In addition, decreasing MOI - which corresponds to decreased barcode insertion probability - will result in fewer cells that are barcoded and thus fewer trackable lineages. As such, it rises the necessity to increase the initial cell pool size, to keep the number of traceable lineages. Consequently, concerning bounded resources and the minimal desired errors of lineages identification, inflation of barcodes’ library or the number of cells may be found redundant concerning the desired goal. In other words, a certain barcode library complexity, together with some specific MOI range, might be sufficient for complying with the observation purposes.

### A. MOI Range

Care must be taken in the experiment design when choosing the MOI parameter. On one hand, for very low MOI – which corresponds to very low barcode insertion probability - most of the cells are eventually unlabeled, and thus their lineages are undetectable. Thus, increasing the MOI results in more cells that are barcode labeled. On the other limit, for a high MOI corresponding to high barcode insertion probability, misidentifying lineages due to dropouts is more probable, hence lowering the MOI brings more accuracy to the identification of measured lineages. In SI App. C for example, we show using both an analytical approach and simulation results, that lineages tagged with a small set of barcodes possess less error in lineages’ identification. This is shown by both simulation and analytical analysis using the binomial model. The compromise between the number of traceable lineages to their accuracy is encapsulated in the counter-effect of the MOI on the total number of correctly measured lineages, see Fig. 2.

**FIG. 2.**
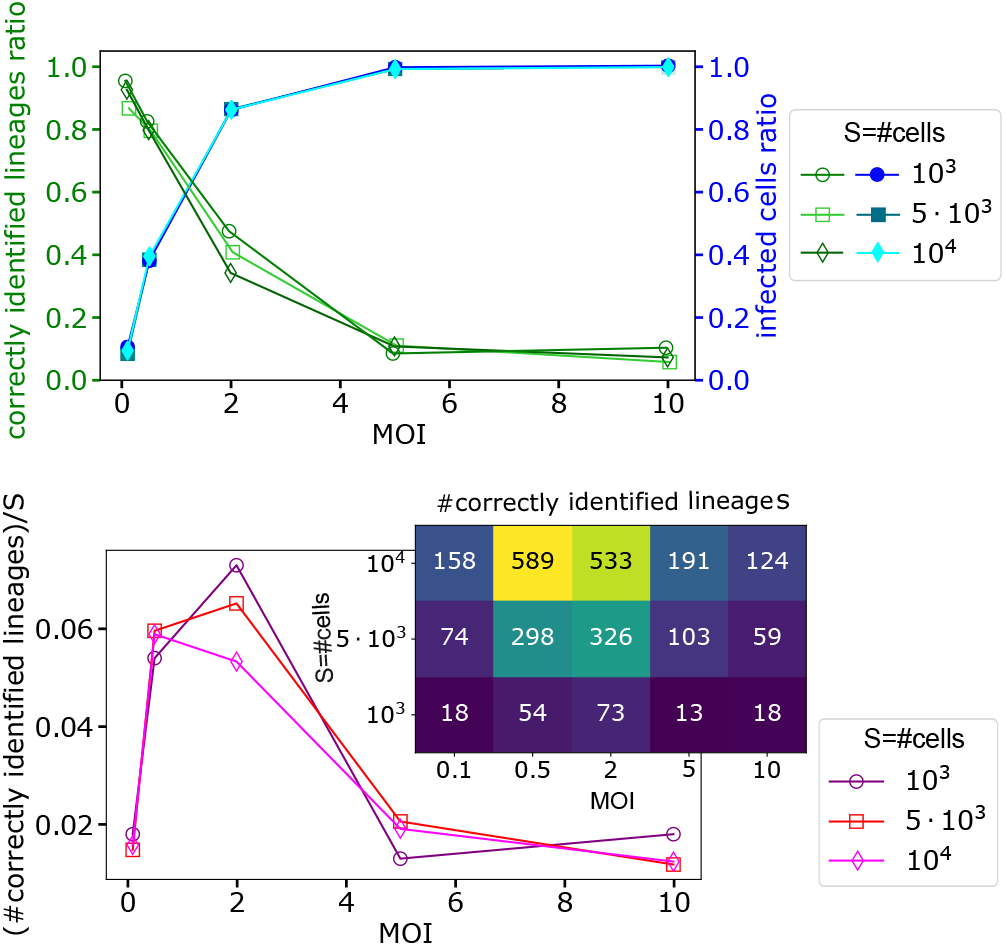
The counter-effect of MOI on the percentages of infected cells and correctly identified lineages. MOI value needs to be chosen carefully, to maximize the number of tracked lineages without significantly damaging the lineages inference quality. Upper panel: Increasing the MOI results in increasing the number of infected cells (blue shades, full markers), while the ratio of correctly identified lineages decreases (green shades, open markers). Lower panel: The number of lineages correctly observed throughout three snapshots; at generations 5, 10, and 15, present non-monotonous behavior. Inset: the number of correctly measured lineages. In here we use the number of cells as *S* = 10^3^, 5·10^3^, 10^4^, and the number of barcodes is 100-fold larger than the initial number of cells with 10% dropouts.

For example, let us consider a given dropout percentage, say 10%, which in reality is given by experimental limitations [21, 22]. In Fig. 2 we show that, as is described above, increasing the MOI has a trade-off; more barcoded cells but lower accuracy of lineages inference. Additionally, the number of correctly observed lineages presents non-monotonous behavior with some range of MOI with a maximal number of perfectly observed lineages. As a result, there is an MOI range that is ‘optimal’ in a sense of bounded resources. The latter refers to the practical limitations of the prepared cells to be infected, and the library complexity. For example, to track correctly at least a given number of lineages, in some scenarios where little less accuracy of lineages’ inference may be tolerated for the specific research goals, it might be easier to increase the MOI from 0.1 to 0.5, than to increase the number of prepared cells of interest, as is demonstrated in Fig. 2. Additionally, in SI App. D we show analytically the non-montonous behavior of the number of correctly identified lineages versus the MOI for the binomial model, which is the qualitatively agrees with the simulation results.

We note that a similar behavior, the obtained trade-off with changing the MOI and thus the concave shape of the number of correctly identified lineages, may be more general and is found for other scenarios; for various percentage dropouts and barcodes library complexity, as well as for non-uniform barcodes and cell distribution. However, for those other situations, simulations together with probabilistic analyses suggest that a different optimal range of MOI may be found. For example, we show in SI Apps. D-F that a low dropout percentage provides better lineages’ identification accuracy, thus higher MOI may be used without significantly diminishing the precision of lineages’ discerning. Worth mentioning that for a very low MOI of 0.1, results appear nearly independent of dropout percentage, as shown in SI App. F. In addition, the ‘optimality’ of MOI may differ with the needed accuracy and even may depend on the choice of the accuracy index. For instance, one may choose MOI considering the correctly clustered *cells* instead of the correctly identified *lineages*. For further discussion and simulation results of the accuracy index in SI App. G.

As a result, quantitatively, the argument MOI which provides maximal correctly identified lineages may vary with properties of the underlying experiment; such as the barcodes pool skewness, the cells susceptibility, cells’ growth homogeneity, the passaging frequency the percentage of the sampled subculture, and low-expression dropouts. Additionally, some experimental purposes may compromise with fewer tracked lineages but gain better accuracy, thus the requirements and goals of the experimental study must be considered as well. Therefore, choosing the MOI must be done with care due to its significant effect on the overall outcomes; both the tracked lineages number and their correctness. Practically, choosing the MOI can be done by simulations, or by using a small experimental ‘sandbox’ system. We note that the number of perfectly measured lineages linearly depends on the number of pre-prepared cells, but the percentage of these correctly measured lineages from all measured ones is independent of the system size, see Fig. 2.

### B. Barcodes Library Complexity

Interestingly, increasing the barcode library complexity beyond some value does not necessarily increase lineages inference quality significantly. Nonetheless, we clarify that a minimal barcode library complexity is needed to guarantee unique barcode sets in the seeding stage and minimize sets overlap. For the example presented in Fig. 3 with a given dropouts percentage of 10%, one can use a not-too-large barcodes library; e.g. *B* ≳ 100*S*, or even *B* ≳ 10*S*, where *S* and *B* are the numbers of pre-prepared cells and barcodes pool complexity respectively, see Fig. 3. That is in agreement with the empirical rule given in [18], that the ratio between cells and barcodes should be 1:100 at least.

**FIG. 3.**
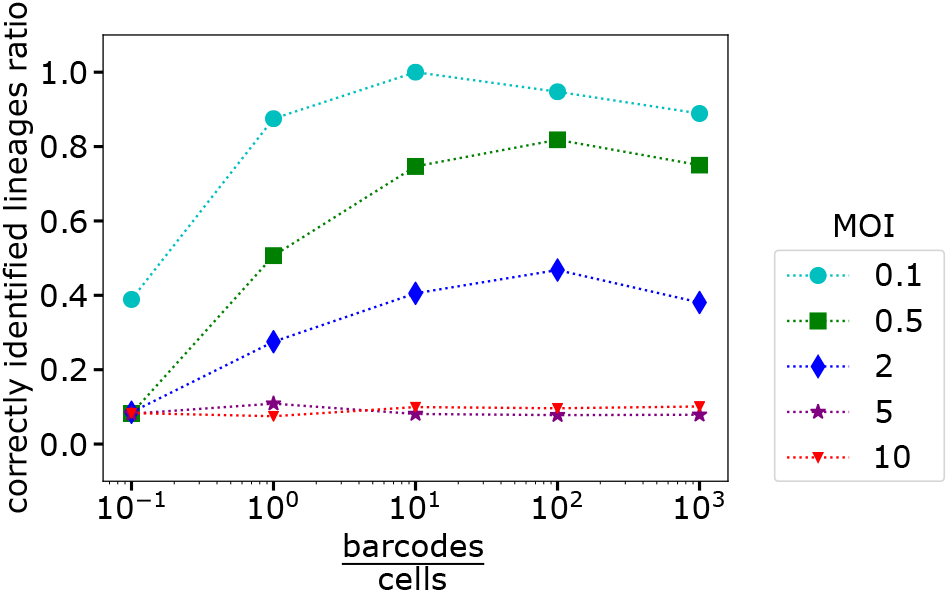
The dependence of barcode library complexity. Simulations suggest that increasing the diversity of potential seeded barcodes does necessarily imply improvements in the lineage tracking quality. Here the number of cells is *S* = 10^3^ with 10% dropouts.

Importantly, for other dropouts’ percentages and barcodes’ frequency skewness, the minimal needed complexity of the barcode pool to guarantee the improbability of sets’ overlap may differ from the above-suggested value. In addition, as was stated above, different experimental requirements may need for other minimal complexity of the barcodes’ pool. For example, as we analytically assess in SI App. H for the binomial model, the MOI itself together with the determined threshold might affect the needed size of the barcodes’ pool. Obviously, for a poor choice of MOI and a high probability of dropouts, even increasing the barcode pool complexity does not sufficiently improve the bad quality of the results, see for example Fig. 3 for MOI=5, 10.

### C. Deviation from Binomial Distribution

Up until now, we have assumed that the barcodes are uniformly distributed, and thus have equal chances to get inserted into a cell. In addition, we have considered all cells are equally susceptible to being barcoded. Nevertheless, in a real experimental scenario that might not be the case. For example, in [9, 27, 28] deviation from the equally distributed barcodes is reported. Still, as briefly mentioned, the above statements are valid even for nonuniform distributed barcodes, where some barcodes are more probable to be delivered to cells. We examine such a scenario; where both cell susceptibility and barcodes’ library are non-uniform. A qualitative similar behavior is obtained; increasing MOI presents a trade-off where more barcoded cells with lower lineages’ tracking quality. The number of correctly inferred lineages presents non-monotonous behavior with a maximal value, see simulation results in App. F.

## IV. DISCUSSION

Careful planning of an experiment is needed to track a given number of cellular lineages using barcodes, while taking into account the preconditions and tunable parameters of the setup such as the barcode insertion probability and the number of cells and barcodes needed, bearing in mind that resources are limited, e.g., the practical number of pre-prepared cells and barcodes is bounded. We have studied the effect of the features of experiments and the importance of carefully choosing the tunable parameters on the quality of the cellular lineages tracking. To do so, we use a ‘null model’ which considers cells’ passaging, barcode insertion, and dropouts. In particular, we examined the effect of the MOI, the barcode pool complexity, the dropout probability, and the deviation from uniformly distributed barcodes and cells.

The underlying features of an experiment have a high impact on the overall quality of lineages’ tracking hence is important to examine them. We have recovered the obvious expected truth that low dropout percentages provide a better quality of lineage tracking - see in SI Apps. C-F. Exceptionally, for MOI=0.1, simulation results appear nearly independent of dropout percentage, as shown in SI App. F - hence such a low MOI seems to be ‘tolerant’ to dropout rate variability - and in that sense might be beneficial. Barcode reading quality depends on the sequencing depth and coverages. A higher sequencing depth generates more informational reads, and thus increases the robustness of scientific findings. Nevertheless, the higher coverage of sequencing inevitably requires higher costs. Therefore, sequencing depth and coverages must be carefully chosen to optimize the experiment setup [36]. In addition in RNA sequencing, the low-expression dropouts rate is practically challenging to reduce in an experimental setup since changing the gene-expression threshold might inflate false-negative and thus affect the validity of reads [22]. Therefore we assume the dropout rate parameter is an outcome of the observations analysis pipeline. The precise dropout percentage is unknown but it can be estimated, e.g., by differential expression signature [21]. Dropout percentages are reported in a range of values - from a few percentages to ∼80% gene dropouts - and may vary among cells, depending on the quality of a particular library, cell type, or RNA-seq protocol. However, we typically find that dropout percentages lie within the interval of 10% to 50% [21, 23, 37–39].

Interestingly, we have found that one may increase the number of propagated lineages, for example by changing the MOI which is a tunable parameter. This parameter presents a counter-effect upon the accomplishment of the multiple-lineages-tracking goal; increasing the MOI provides more lineages to be tracked, but damages the quality of lineages identification thus fewer lineages are correctly measured. Decreasing the MOI provides only several trackable lineages but most are correctly identified. The balance between these two effects yields an ‘optimal’ range of MOI which gives the maximal number of correctly identified propagated lineages under a minimal accuracy needed.

The ensemble size of prepared targeted cells has also a significant impact on the overall accomplishment of the experiment goals. Where the number of barcoded cells and the number of correctly measured lineages are both linearly dependent on the number of pre-prepared cells (see Fig. 2), where the inherent randomness needs to be considered as well. However, expanding the cells’ ensemble may be technically hard, and beyond some ensemble sizes even impractical.

Thus, with a given ensemble size of prepared cells, we have presented a limitation over the number of correctly identified lineages - where we found that a maximum of ∼7% of prepared cells are eventually properly identified with their lineages. As mentioned, correctly identified lineages are from cells that are labeled with barcodes, resampled in each passaging, and correctly observed throughout the timeframes. We note, that each step; labeling, resampling, and observing, might reduce significantly the percentage of overall tracked lineages. For example, in MOI=2, around 0.86 cells are labeled at initializing step. Then our simulation yields that only 0.187 labeled cells at t=0 are resampled throughout all passaging. Thus only around 16% of prepared cells are labeled and propagated - even when the observation of barcodes is perfectly executed without any dropouts or other measurement errors (see SI App. B). The passaging itself with its properties - its frequency and the sampled subpopulation ratio - is governed by neutral dynamics where all the cells possess the same growth rate. Controlling the frequency and ratio of passaging might also be beneficial to lineages tracking task.

Accordingly, we have suggested that a careful choice of MOI may be beneficial. For example let’s assume that in some experimental scenarios, one aims to measure at least 100 lineages and may ‘tolerate’ that at least 70% from measured lineages is correctly inferred. Then, in Fig. 2, we demonstrate that one may choose to prepare 10^4^ cells with planned MOI=0.1. However, if such an initial number of cells is hard or even impossible to achieve, one may compromise the accuracy, such that preparing only 5 ·10^3^ cells with MOI=0.5 might be sufficient for some specific research goal.

To clarify, research goals and limitations may vary from one to another, together with other experiment properties that are different from our model, thus our simulated results do not determine any specific value as the gold standard. In addition, other lineages-inference-quality scorings may be used, e.g., using the number of *cells* correctly associated with their lineage, instead of the number of non-mixed measured *lineages*. Using other labeling scores may result in varied consequences, even when yielded from the same data, see further discussion in App. G. Yet, we capture the importance of cautious planning of an experiment with all its tunable parameters, similarly as qualitatively demonstrated in our simulation results.

As mentioned, from a practical perspective choosing the MOI can be done by simulations, or by using a small experimental ‘sandbox’ system. To do so, we note that the system size *S* linearly affects some quantities, e.g. the absolute number of labeled cells, the number of measured lineages, and the number of correctly measured lineages. Conversely, some values are independent of the system size, such as the percentage of labeled cells and the ratio between the number of correctly measured lineages to all observed lineages.

Moreover, we have assessed the effect of barcode pool complexity on the lineages’ tracking quality. We have found that beyond a given complexity of the barcodes’ pool, there is no advantage to increasing its complexity further since it does not contribute to the lineages inference task. We note though, that a minimal barcode library complexity is needed to reduce the probability of overlap between barcodes’ sets.

## ACKNOWLEDGMENTS

The authors acknowledge MbD and NSERC Discovery for funding support. We thank Sophie McGibbon-Gardner for the discussions and comments.

