## Supplementary Information for "Limitations and Optimizations of Cellular Lineages Tracking"

#### Appendix A: Uniform Insertions and Dropouts - Analytic and Simulation Results

##### 1. Uniform Insertion

Assume that the probability for a barcode to be inserted into a cell is independent of barcode identity, the cell's identity, or any cell's state (e.g., susceptibility to be infected due to prior infections). In that case, the probability of any barcode to be inserted is simply a constant $p_i$ [the subscript $(.)_i$ refers to 'insertion']. In that case, the insertion is binomial, where the probability to have a cell with a barcodes-set size of $L$ is given by

$$P(L) = \text{Binom}(B, p_i) = \binom{B}{L} p_i^L (1 - p_i)^{B-L} \quad (\text{A1})$$

where $B$ is the diversity (also refers to as complexity) of the barcodes' pool. As known, since the insertion probability of a single barcode is very low, i.e., $p_i \ll 1$, and the complexity of the barcodes pool is very high $B \gg 1$, the size of the barcodes' set is approximately Poisson [1] namely,

$$P(L) \approx \text{Poisson}(M) = \frac{e^{-M} M^L}{L!} \quad (\text{A2})$$

where $\langle L \rangle = M \equiv p_i B$ is the MOI commonly described in the literature. Here, the mean number of infected cells is simply $1 - e^{-M}$. Note that through the simulation analysis and visualization, $p_i$ is determined by the barcode pool diversity $B$ and the MOI $M$. Details about the binomial and Poisson distributions may be found in [1].

##### 2. Uniform Dropouts

The binomial dropout is based on the same assumptions as binomial insertion; no barcode is more probable to be dropped and no cell is more probable to lose its barcodes. Therefore, the *observed* distribution of barcodes-set size, which takes into account both insertion and dropout is

$$\text{PDF}(L) = \text{Binom}[B, p_i(1 - p_d)] = \binom{B}{L} [p_i(1 - p_d)]^L [1 - p_i(1 - p_d)]^{B-L} \quad (\text{A3})$$

where, as mentioned above, the *measured* MOI is given by $\langle L \rangle = M = B p_i (1 - p_d)$.

As mentioned, in that simple scenario both barcodes and cells are uniformly distributed, in the sense that they are functionally identical; all cells are equally susceptible, and all barcodes have the same probability to be inserted. Therefore, the number of cells with a given barcode also follows binomial distribution;

$$\text{PDF}(C) = \text{Binom}[S, p_i(1 - p_d)] = \binom{S}{C} [p_i(1 - p_d)]^C [1 - p_i(1 - p_d)]^{S-C} \quad (\text{A4})$$

where $C$ is the random variable that represents the number of cells with a given barcode, $S$ is the number of cells potentially seeded in the initial state, and $p_i$ and $p_d$ are the probability of a single barcode to be inserted and dropped. Here $\langle C \rangle = Sp_i(1 - p_d) = S\langle L \rangle/B = (S/B) \cdot M$, which is linear with $M$.

In Fig. 1 we present the statistical features of the binomial case, in agreement with the predicted values from the binomial model provided in Eqs. (A3) and (A4).

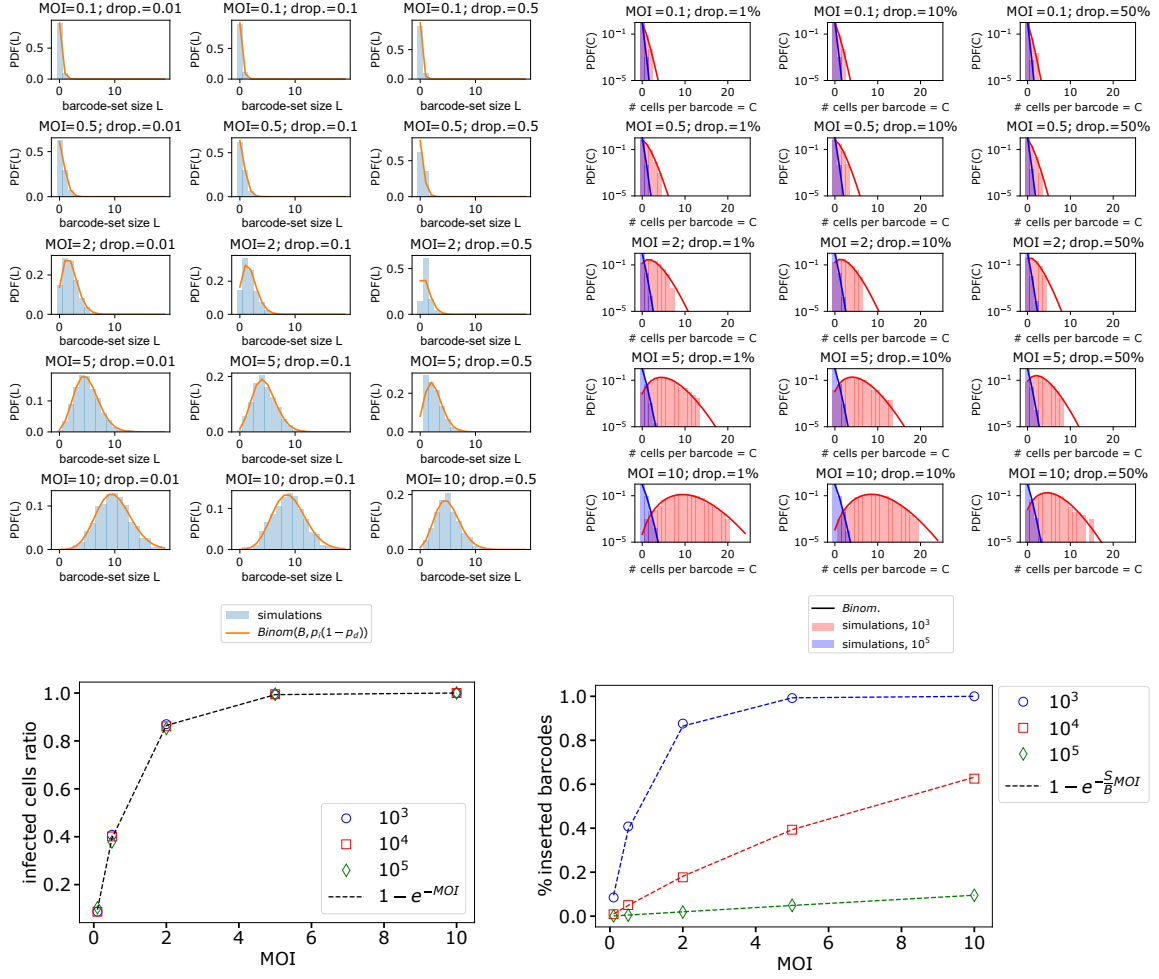

FIG. 1. Statistical features of the uniform insertion. Upper left panel: Measured barcodes-set size distribution $PDF(L)$ for various MOI and dropout probability follows the binomial distribution. Here we present the results for barcodes' pool diversity of $10^5$ barcodes. Upper right panel: Number of cells that are infected by a given barcode; $PDF(C)$. We present barcode pool diversity of $10^3$ (red) and $10^5$ (blue). The solid curves are their corresponding binomial distributions. Lower left panel: the fraction of infected cells, without any dropouts, increases with the MOI and is independent of the barcodes' pool diversity. Lower right panel: the fraction of barcodes inserted (with no dropouts) in the seeded stage, with its dependence on the MOI and the barcodes pool diversity, see details in the main text.

### Appendix B: Time Propagation

Up until now, we described the seeding procedure - which aims to associate each cell with a set of barcodes. The following step is to use only labeled cells and remove the unlabeled ones (e.g. by using puromycin selection). Thus, at that point, we have what we call the ensemble of labeled cells in the initial time $t = 0$. Then, the ensemble of labeled cells is multiplied (such that every mother cell divides into exactly two daughter cells) and sampled such that the size of the ensemble of cells returns to its original size from the initial stage, a procedure that is called passaging [2]. We repeat the division-sampling process 15 times, which corresponds to 15 generations. Note that, as mentioned in the main text, the majority of lineages are lost during the time propagation. In our simulation, we have found that around 18.7% of initial cells are propagated and resampled through the 15 generations. That fraction is given from simulation results shown in Fig. 2, where we present the histogram for several repetitions.

In generations 5, 10, and 15 we keep for analysis the data of the ensemble of barcodes' cassettes corresponding to the labeled cells. Hitherto we generated synthetic data of cellular lineages in four snapshots; at generations 0, 5, 10, and 15. This represents the ground truth of the propagated lineages and is used to examine our results. Additionally, in each time frame; 0, 5, 10, and 15, to capture the difficulty of a realistic experimental observation we apply our dropout modeling too - which means that some of the barcodes dropped and were not observed. We illustrate our model in Fig. 3

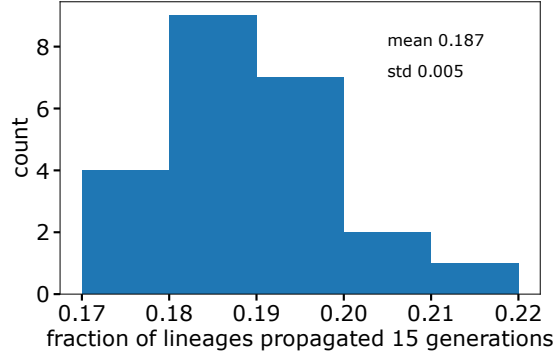

FIG. 2. The number of lineages observed (without dropouts) after generation 15 compared with the initial number of clones. Most of the lineages are lost during the resampling procedure. As a result, even with a perfect observation without any noise emerging from dropouts, on average only 18.7% of the initially labeled cells are propagated and resampled.

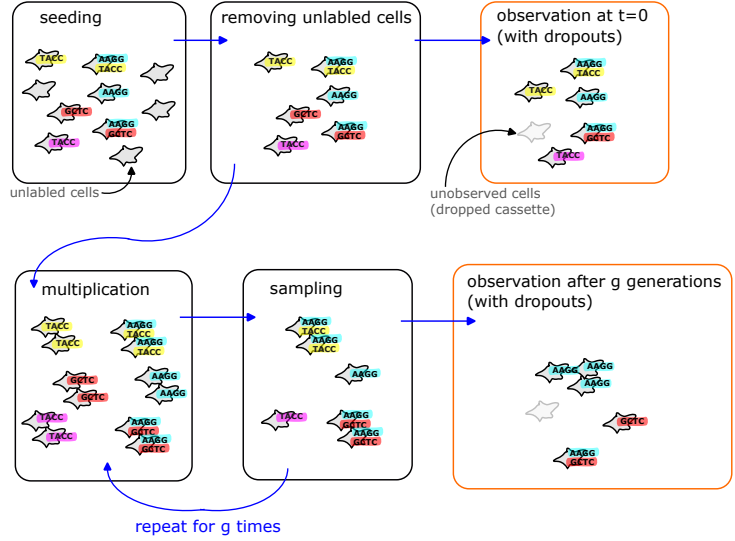

FIG. 3. Illustration of our synthetic data generation procedure. We start with seeding - the insertion of barcodes into cells. As described in the text, some cassettes have more than one barcode, and some cells remain unlabeled. These latter are removed from the cells, thus all cells under examination are labeled with barcodes' cassettes. In each generation, we assume cell division and sampling (also known as passaging). In the seeding stage, and after $g$ generation, we keep the cells' barcode data - both the true propagated lineages (without dropouts) and with dropouts to mimic realistic observations.

### Appendix C: Dependence on Barcodes-Set Size - Additional Simulation Results and Exact Analysis from the Uniform Insertion and Dropout Model

Large sets of barcodes are troublesome for lineages recording due to the dropouts present. There, the possibility of misreading of barcodes' sets is higher and thus leads to a wrong inference of a lineage, even when the initial barcodes' sets that were seeded are unique.

As mentioned, in our model we assume the dropouts occur binomially. Thus, the probability that two barcodes' sets, conditioned to be identical where no dropouts occur and assumed to be with size $L$, are observed identically where considering dropout, is given by

$$\begin{aligned} \text{Prob}(\text{barcodes in cell 1} = \text{barcodes in cell 2} | L) &= \\ &= \sum_{l=0}^{L-1} \binom{L}{l} p_d^{2l} (1 - p_d)^{2L-2l} = (1 - p_d)^{2L} \left( \frac{2p_d^2 - 2p_d + 1}{(p_d - 1)^2} \right)^L - p_d^{2L}. \end{aligned} \quad (\text{C1})$$

In Fig. 4, we plot the above mathematical expression, with the dropout probabilities $p_d = 0.01, 0.1, 0.5$ and barcodes-set sizes $L$ are integers spanning between 1 to 10.

Interestingly, this probability is a nonmonotonic function neither with $L$ nor with $p_d$. For example, for a large dropout probability, say $2/3 < p_d < 1$, cells with a set of two barcodes are more probable to be measured identically when compared to cassettes of a single barcode. Therefore for such high dropout probabilities, planning the experiment to insert longer cassettes might be beneficial. Nevertheless, we emphasize that high dropout probabilities must be avoided if possible.

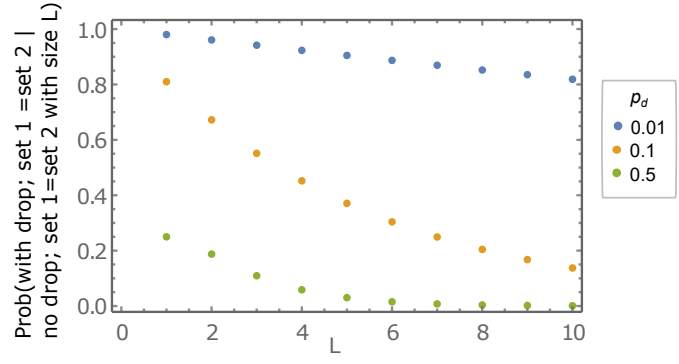

FIG. 4. The probability that two barcodes' sets are observed identically, conditionally their true sets are identical and with a size of $L$, as given in Eq. C1 where note that $L$ is an integer.

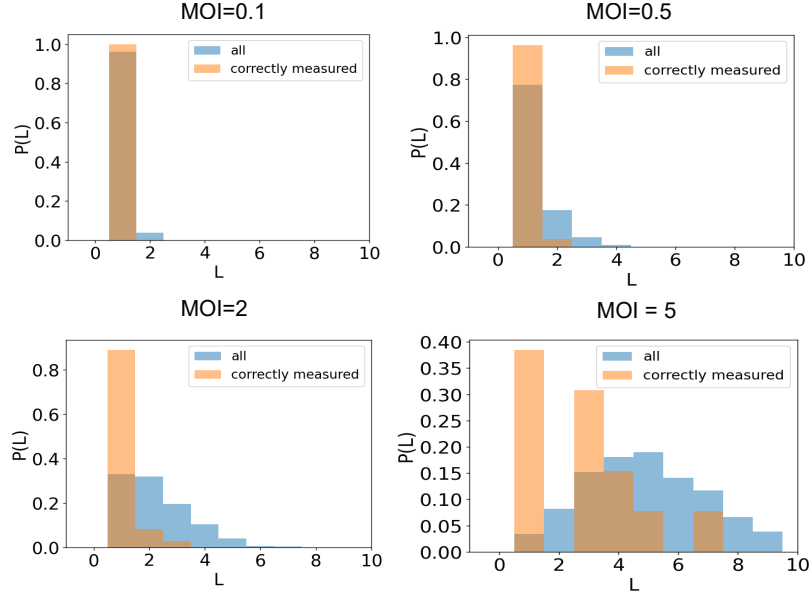

FIG. 5. Correctly measured lineages are mostly labeled with small sets of barcodes. Here we present our simulation results for an ensemble of $S = 10^3$ cells, with barcode pool complexity of $B = 10^5$ barcodes and 10% dropouts. The distribution of barcodes' set sizes of *labeled* cells follows the truncated Poisson probability density function (removing the unlabeled cells, the ones with $L = 0$), and are represented by blue histograms corresponding to the MOI (see also Fig. 1 for the comparison with Poisson). The eventually perfectly correctly measured lineages have barcodes' set size distribution with a mode at $L = 1$ as shown by the orange histograms.

##### Appendix D: Dependence on the MOI - Exact Analysis from the Uniform Insertion and Dropout Model

In the simulation results shown in the main text, we present the number of perfectly inferred lineages. We consider perfectly measured lineages if all its inferred associated cells are truly from that lineage, and all its measured cells are inferred to be from that lineage, namely non-mixed lineages.

Consider that cells are associated with a given lineage if and only if their *measured* barcodes' sets are identical. Then the estimation of the number of lineages that are measured correctly is approximately

$$\#(\text{perfect lineages}) = \# \text{cells} \times \text{Prob}(\text{propagated}) \times \text{Prob}(\text{identically measured cassettes}). \quad (\text{D1})$$

Here, the probability that two cells are measured with the same set of barcodes, given that the true (without dropout) barcodes-set size is $L$, is given in Eq. (C1). As expected, smaller sets of barcodes and low dropout probability increase the probability that the two cells have identically measured sets of barcodes. For the following illustration, we consider that the probability that a set has a size of $L$ is binomial, and approximately a Poisson distribution as discussed above. Therefore, the probability to measure two identical sets is

$$\begin{aligned} \text{Prob}(\text{measure two identical sets}) = & \quad (\text{D2}) \\ \sum_L \text{Prob}(\text{barcodes in cell 1} = \text{barcodes in cell 2} | L) P(L) \end{aligned}$$

which can be calculated numerically, as is given in Fig. 6. As is shown, for very low MOI, many cells are unlabeled and thus unmeasured, hence the very low probability. On the other extreme, for very high MOI, dropout may cause different sets to be measured hence measuring two identical sets of barcodes becomes less probable. Importantly, the probability to measure two cells with an identical barcode cassette yields a concave function with some maximum point which depends on the dropout probability. Interestingly, the argument of the maxima decreases with dropout probability.

To estimate the number of perfect measured lineages as the given simulation results in the main text, one needs to take into account three additional variables; First, $N$ with its variability, where $N$ is the number of *cells* within a given lineage across the measured time frames. Generalizing the above calculation for the probability of measuring $N$ identical cassettes is straightforward, and the qualitative behavior given in Fig. 6 holds, however, $N$ is unknown. Second, the number of perfect inferred lineages depends on the probability of an initial lineage to be propagated until generation 15 without getting lost in passaging (sampling) steps. Our simulation suggests 18.7% of initially labeled cells are propagated throughout all 15 generations (see Fig. 2). The third and last component is given from the following. In our simulation, we infer lineages even when the sets of barcodes are not perfectly identical, a fact that needs to be taken into the account as well. These three missing elements in our analysis can be taken from the simulated synthetic data, and beyond the scope of this analysis.

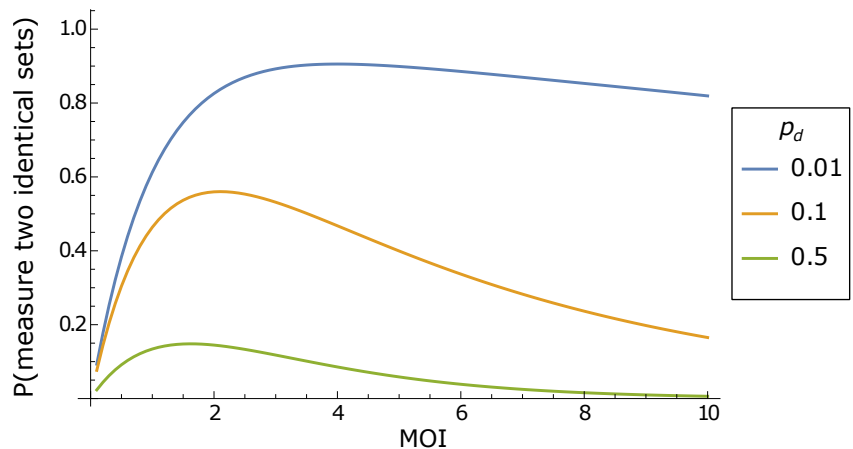

FIG. 6. The probability to measure two cells with identical sets of barcodes. Here we present the analytical expression given in Eq. (D2). For very low MOI, dropout causes cells to be unmeasured and thus the very low values for $\text{MOI} \rightarrow 0$. On the other extreme, for very high MOI, dropout may cause different barcodes' sets to be measured.

### Appendix E: Over-Dispersed Scenario

As mentioned in the main text, in some experimental scenarios, some barcodes can be more abundant than other, and some cells can be more susceptible than other. For example, in some barcode pool preparation procedures, some deviation from uniform barcode abundances toward exponential distribution is reported, see for example in [3, 4].

The index of dispersion $D = \sigma^2/\mu$, also known as the Fano factor or variance to mean ratio (VMR), captures the deviation from Poisson statistics. In the above binomial cases, which can be approximated as Poisson, $D \lesssim 1$. An equality $D = 1$ is expected for pure Poisson distribution. However, for the over-disperse scenario presented here, $D$ may obtain larger values, see simulation results in Fig. 7. For sufficient low MOI values, both Poisson and Exponential models present $D \lesssim 1$.

#### 1. Over-Dispersed Barcodes-Set Size Distribution

In our simulated over-dispersed cases we assume both deviate from binomial; the number of cells with a given barcode, and the number of barcodes within a cell. We choose the size of barcodes' set $L$, to be distributed following the exponential distribution with scaling of $\langle L \rangle$ similar to the MOI of the uniform case. In Fig. 8 we show simulation results of $\text{PDF}(L)$ for the over-dispersed case. We note that in the conventional exponential distribution, the random variable is continuous, whereas in our case, the number of inserted barcodes is discrete. Hence, the random variable $L$ is chosen from the exponential distribution with rounded values. In other words, the random set size is given by $L = \text{round}[\hat{L}]$ with $\hat{L} \sim \text{Exp}(\text{MOI})$, see Fig. 8.

#### 2. Over-Dispersed Cells per Barcode Distribution

As mentioned, we consider both cells and barcodes to be non-uniform. Here we include the case where, beyond some cells being more susceptible, some barcodes are more likely to be inserted. In the synthetic generated data presented here, we choose the probability for the $n$-th barcode, $b_n$, to be inserted as $p_i(b_n) \propto \exp(-10n/B)$, where $n$ is some arbitrary (but fixed) index of barcodes, and subscript $(.)_i$ refers to insertion. Note that the ratio of probabilities to be chosen between the least and most probable barcodes is $e^{-10}$, in comparison with the uniform case where all barcodes can be chosen equally. To guarantee the exponential size of barcodes' set with a known mean (we aim to generate $\langle L \rangle = \text{MOI}$), our synthetic data is generated as follows. For a given cell, we randomize the set size $L$ following some $\text{PDF}(L)$. Then, we 'fill' the set with barcodes chosen randomly considering $p_i(b_n)$. If the randomly chosen barcode already exists in the cassette we choose another one, until the set size reaches $L$. Note that it yields that the number of cells with a given barcode presents a distribution that is heavier than exponential, see simulation results in Fig. 8.

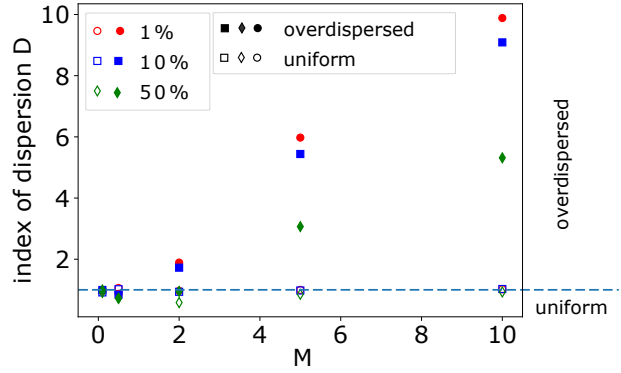

FIG. 7. Index of dispersion $D$ for barcodes-set size distribution $\text{PDF}(L)$, for the generated synthetic data used in our analysis. The open markers represent data from the uniform cells and barcodes, thus $D$ is expected to be close to one. For our over-dispersed generated data, $D$ yields high values for large $M$. Here $M = Bp_i$ is the ideal MOI, with no dropouts, for the uniform identical cells and barcodes, and $M = \langle L \rangle$ for the exponentially distributed barcode-set size without dropouts events. The dashed line is drawn at $D = 1$ and represents the transition between the scenarios. Here we present the results for barcode pool diversity of $10^5$ barcodes.

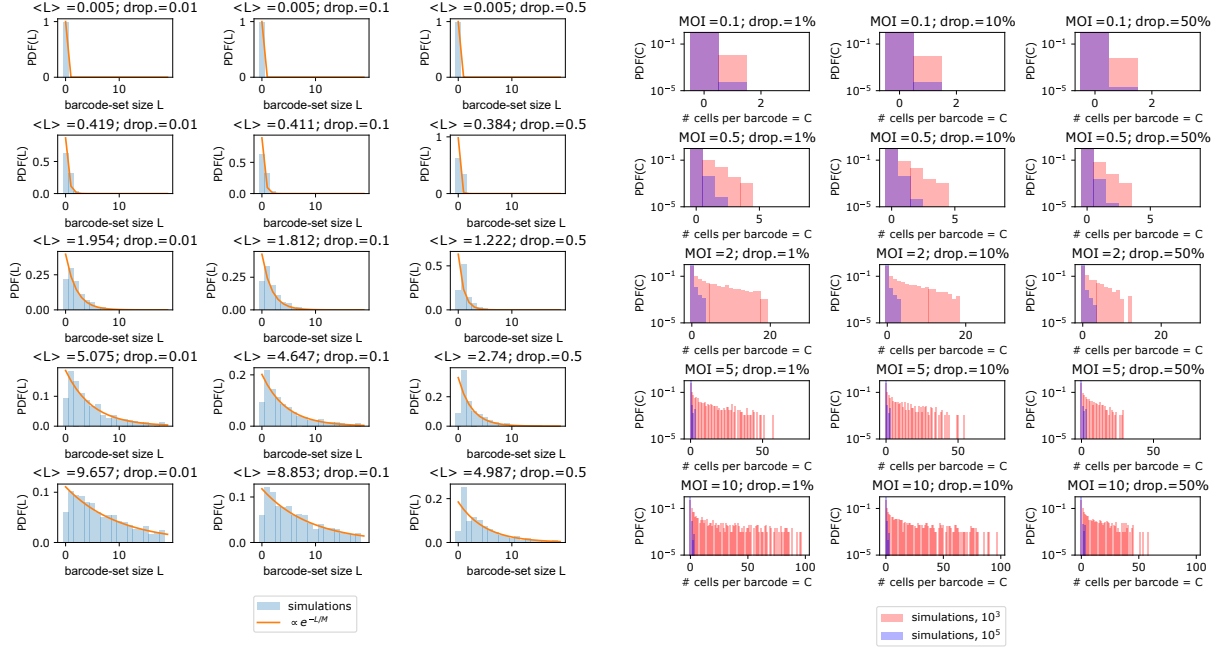

FIG. 8. Statistical features of the over-dispersed scenario. Left panel: Measured barcodes-set size distribution. In this simulation, the barcodes-set size distribution follows exponential distribution; $\propto \exp(-L/M)$ where its scale $M = \langle L \rangle$ aims to be close to the MOI presented in the binomial case presented above. Here we present the results for barcode pool diversity of $10^5$ barcodes. Right panel: The frequency of cells with a given barcode where the barcodes pool diversity is $10^3$ (red) and $10^5$ (blue).

### **Appendix F: Additional Simulation Results**

#### **1. Simulation Results for Over-Dispersed System**

As was mentioned, in some empirical observations a deviation from binomial distribution is found, which might suggest that some cells were more susceptible than others or that some barcodes were more abundant within the pool. However, a similar qualitative behavior - a concave shape of the number of correct lineages - is obtained.

As shown in Fig. 9 the nonmonotonic MOI effect on the number of correctly measured lineages is found in both scenarios; uniform and non-uniform cells and barcodes which correspond to binomial and over-dispersed distributions respectively. Additionally, this qualitative shape is maintained also for low (1%) and high (50%) dropout percentages.

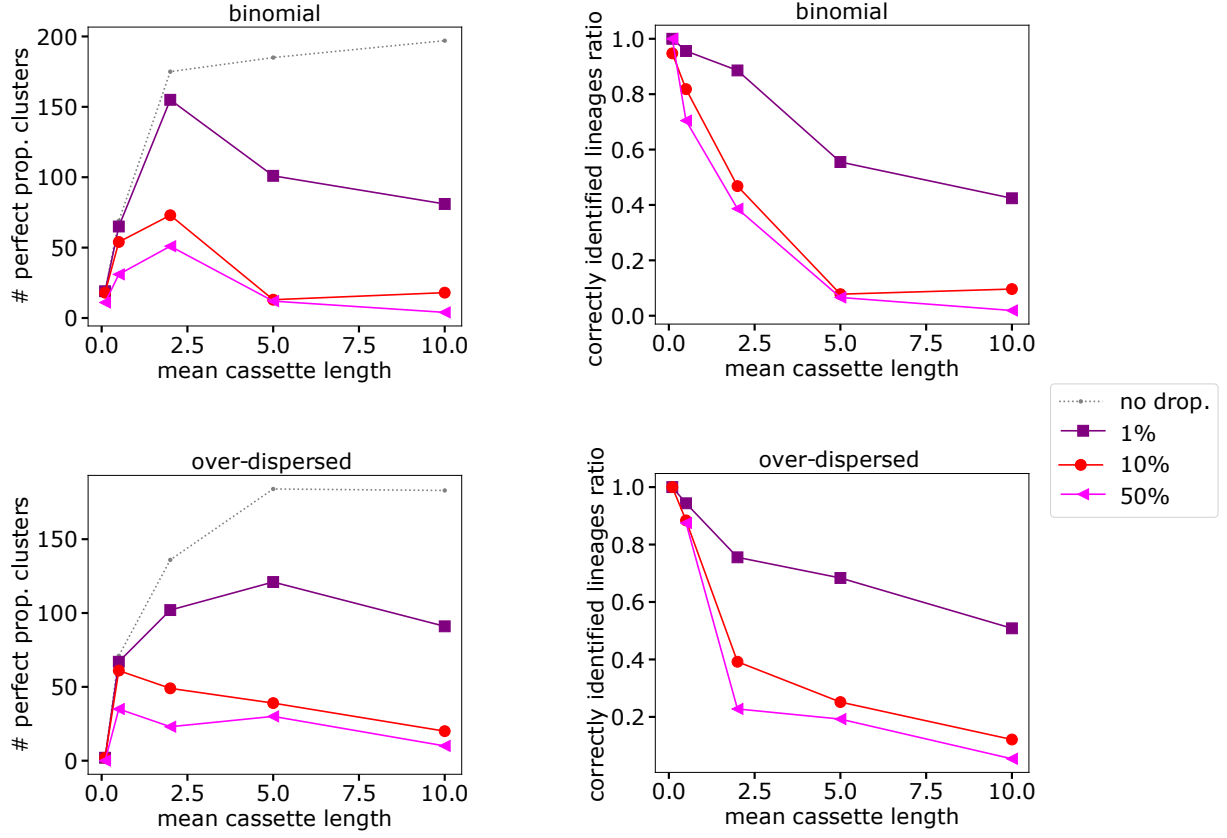

FIG. 9. The number of lineages correctly observed throughout propagation, and the correctly-identified-lineages ratio, for different insertion scenarios and dropout percentages. Upper row; all cells and barcodes are functionally identical, thus the insertions follow binomial statistics. Lower row; barcodes' set size is exponentially distributed, barcode library is skewed (see text). Here we use the number of cells = $10^3$, and the number of barcodes = $10^5$ for the simulation.

### Appendix G: Simulations Analysis

To recover the lineages, while considering dropouts, one needs to develop a clustering method, which associates the observed barcodes in cells, with their appropriate lineages, even when cells do not share the exact overlapping barcodes cassette. In the following, we discuss thoroughly our clustering building procedure. Afterward, we examine the clustering quality concerning different aspects relevant to experimental plans.

#### 1. Overlapping Percentages Metric

To cluster together cells corresponding to their barcode cassette, one needs to consider some metric, or an “effective” distance, between two sets of barcodes, which somewhat captures the plausibility that the two cassettes are from the same lineage. Thus, the distance between two sets is small when it is plausible that the two sets are from the same lineage, and large if the two cassettes are unlikely to be from the same lineage.

Assume that the cross-affinity between two barcode sets is determined by the overlapping percentage between the two sets. There, two barcode sets are from the same lineage if their sets share many barcodes, i.e., where they have a large percentage overlap. The metric, known as Jaccard distance, is defined as follows;

$$d(X, Y) \equiv 1 - \frac{|X \cap Y|}{|X \cup Y|} \quad (\text{G1})$$

where $X$ and $Y$ are two barcode sets, and $|\cdot|$ refers to set’s cardinality. Note that, as desired, for a perfect overlap between the sets, $d = 0$ such that these sets must belong to the same lineage. On the other extreme, for two disjoint sets, $d = 1$ such that they are unlikely to be from the same lineage.

#### 2. Threshold

After the construction of the distance matrix between every pair of barcode sets, we aim here to determine the threshold of pairing. The latter refers to a given maximal distance between two barcode sets to consider them with the same lineage.

Choosing the threshold might have a great impact on the overall quality of lineage identification. On the one hand, too large a threshold results in the wrong association of cells with their true lineage, where the clustering homogeneity is poor and lineages are wrongly merged. On the other hand, too small a threshold, lead to many cells which are wrongly considered as different lineages, but they are truly from the same lineage, namely bad clusters completeness, and lineages are wrongly split. The threshold was chosen in such a way that it in-balanced the two types of clustering errors - both merging and splitting of lineages - as follows.

#### 3. Goodness of Clustering

In the main text, we present in Fig. 2 the quality of the identification of lineages - which is considered there as the ratio between the number of correctly identified lineages to the number of all identified lineages. That value, for various MOI and dropout percentages, is also shown in Fig. 10 upper panel. That ratio captures the fraction of correctly identified *lineages*, but not how many *cells* are identified correctly with their true lineages, thus that score might yield very low values. For example, a lineage does not consider as correctly identified if one (or more) of the cells from that lineage is wrongly associated with another lineage or if one cell from another lineage is deduced to be from that lineage. Therefore, in the following, we also discuss two other clustering scores which capture the lineages' identification quality at the individual cells' level.

Generally, one can use various quantities to capture the overall clustering quality, where each may have advantages and disadvantages. We note that other clustering scores might be relevant to the experimental scenario and its purposes. The Fowlkes-Mallows (FM) index is defined as the geometric mean between precision and sensitivity. Thus, it captures both the true-positive and positive-predictive rates of the cells. FM index yield value of 1 where the clusters (=lineages) are perfectly inferred, and FM=0 for random classification of cells to lineages.

Conventionally, the errors in lineage construction may be from two types. One is where cells from different lineages are clustered together to be in the same lineage, namely lineages' merging errors. The other one is when cells from the same lineage are identified as two different lineages, what we called splitting errors. The two features of clustering quality can be quantified using homogeneity and completeness measures. Homogeneity measures how much the cells in a given cluster are truly from the same lineage, and thus captures the merging errors. Completeness, on the other hand, measures how many cells that are from the same lineage are classified as the same cluster, hence quantifying the splitting errors. The v-measure captures the arithmetic average between the homogeneity and completeness scores. This quantity presents a single-value measure for the quality of the clustering. The v-measure is a scalar number in $[0, 1]$ where 0 is found for bad clustering and 1 is given for perfect clusters. However, The v-measure random labeling won't yield zero scores especially when the number of clusters is large.

In Fig. 10 we show the lineages construction scores; correctly identified lineages ratio, FM score, and V-measure present qualitatively similar behavior - higher MOI or higher dropout percentages yield less accuracy.

##### **4. Number of Correctly Identified Lineages**

To achieve the main purpose of the barcoding procedure, and to gain insights into the overall cellular dynamics, solely clustering cells into clones in a single snapshot is insufficient. The dynamical features are captured in several snapshots and their analysis, where the clones are expected to be detected throughout these time frames, thus the lineages are inferred.

To quantify the achievement of this goal, we count how many lineages are perfectly identified throughout three-time frames of the simulation; generations 5, 10, and 15. A lineage is considered perfect if all cells concluded to be from that lineage are truly from that lineage, and no cell from that lineage is wrongly associated with other lineages. Therefore, the correctly identified lineages ratio is given by the percentage of correctly identified lineages from the total inferred lineages.

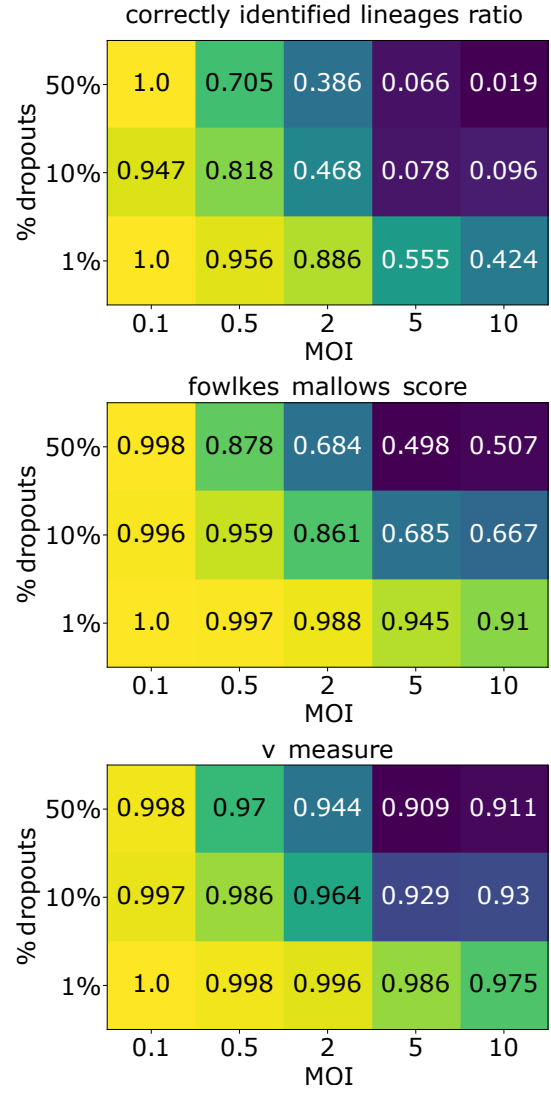

FIG. 10. Lineages identification scores. The upper panel shows the ratio between the number of correctly measured lineages to the number of all propagated lineages. [Note that the results for 10% are also given in the main text, but there we round to 2 decimal places in contrast to here where 3 decimal places are shown.] For the same identified lineages with the same data, we present another two clustering scores; Fowlkes-Mallows score (middle panel) and v-measure (lower panel).

### Appendix H: Minimal Barcodes' Pool Complexity - Exact Analysis from the Uniform Insertion and Dropout Model

A highly complex barcode pool is needed for preventing the integration of barcodes more than once. However, as claimed in the main text, beyond some value there is no advantage to increasing the complexity of the barcode pool. To illustrate that statement we assume the Poisson scenario - where both barcodes and cells are functionally identical and thus follow Poisson statistics. The probability that a barcode is inserted more than once is given by

$$\text{Prob}[\text{overlap}] = \sum_{C=2}^S \text{PDF}(C) = \sum_{C=2}^S \text{Binom}[S, p] = 1 - (1 - p)^S - Sp(1 - p)^S \quad (\text{H1})$$

where the parameter $S$ represents the number of cells. $p = p_i(1 - p_d)$ is the *effective* insertion probability that is given by the true insertion and dropout probability $p_i$ and $p_d$ respectively. The effective insertion probability is determined by the MOI and the barcodes' pool complexity as $p = \langle L \rangle / B = \text{MOI} / B$.

The needed barcodes' complexity $B$ is determined by solving $\text{Prob}[\text{overlap}] < \text{Threshold}$, where the threshold is determined by the experimental requirements. Fig. 11 presents the results for $B/S$ for $\text{MOI}=1$, where we examine two thresholds. The first is $\text{Threshold} = 1/S$ which means no more than one barcode is integrated more than once. The other threshold is 0.001 which refers to less than 0.1% of barcodes being integrated more than once. We have found that for $\text{Threshold}=1/S$ the needed barcode complexity is $B \sim S^{3/2}$, while for $\text{Threshold} = 0.001$ $B/S$ is constant, see analytical results in Fig. 11. Our analytical results agree with the simulation results for $S = 10^3$ presented in Fig. ?? shown in the main text.

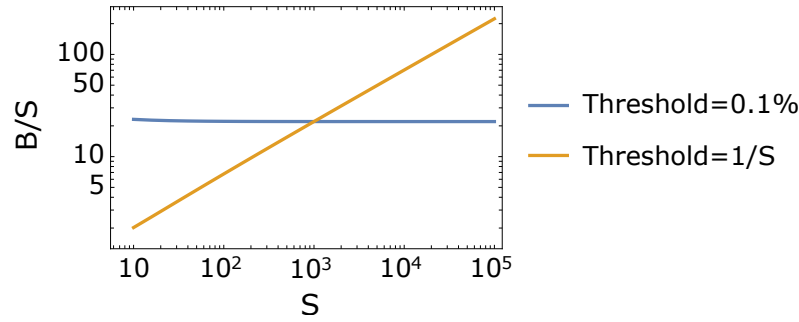

FIG. 11. The barcodes to cells ratio ($B/S$) depends on the experiment requirement. The first is for threshold=0.1% and refers to the case where the probability that two cassettes share the same barcode is less than 0.001. The other one is given by Thershold=1/S and considers the probability that less than one cell shares the same barcodes, meaning that the probability is less than 1/S. Here, the analytical expression for $B/S$ versus the number of cells $S$ is provided by solving Eq. (H1), and is plotted for MOI=1 for the two Threshold requirements (see legend).

- [1] V. K. Rohatgi and A. M. E. Saleh, *An introduction to probability and statistics* (John Wiley & Sons, 2015).
- [2] J. R. Masters and G. N. Stacey, Changing medium and passaging cell lines, *Nature protocols* **2**, 2276 (2007).
- [3] S. N. Porter, L. C. Baker, D. Mittelman, and M. H. Porteus, Lentiviral and targeted cellular barcoding reveals ongoing clonal dynamics of cell lines in vitro and in vivo, *Genome biology* **15**, 1 (2014).
- [4] N. Shakiba, A. Fahmy, G. Jayakumaran, S. McGibbon, L. David, D. Trcka, J. Elbaz, M. C. Puri, A. Nagy, D. van der Kooy, *et al.*, Cell competition during reprogramming gives rise to dominant clones, *Science* **364**, eaan0925 (2019).
